# Social Reactivation of Fear Engrams Enhances Memory Recall

**DOI:** 10.1101/2021.01.07.425728

**Authors:** Abby Basya Finkelstein, Héloïse Leblanc, Rebecca H. Cole, Troy Gallerani, Anahita Vieira, Yosif Zaki, Steve Ramirez

**Affiliations:** Department of Psychological and Brain Sciences, Boston University, Boston, MA, USA; Department of Molecular and Cellular Biology, Harvard University, Cambridge, MA, USA; Boston University School of Medicine, Boston, MA; The Broad Institute of MIT and Harvard, Cambridge, MA, USA; Icahn School of Medicine at Mount Sinai, NYC, NY, USA

**Keywords:** social, c-FOS, memory, engram, dentate gyrus

## Abstract

For group-living species such as humans and rodents, conspecific interactions pervasively shape emotion (1–3), attention (4), and cognitive ability (5–8). Higher-order cognitive processes such as memory within a social brain are thus interlaced with social influences. Traditional laboratory rodent cages offer a limited but nonetheless rich multi-modal landscape of communication, including auditory calls (9–12), chemical signaling (13, 14), and tactile stimulation (15, 16). The absence of such social encounters in singly housed animals results in cognitive impairments and depression-like phenotypes (17), likely obscuring how the social brain has evolved to function. It is thus important to understand the relationship between social context and how individuals process memories. As social interaction recruits hippocampal (18) and amygdalar (19) circuitry that also serves as hubs for non-social memory traces(20–24), we hypothesized that pre-existing ensembles in these regions can be modulated by social experiences and lead to changes in memory expression. Here we show that stressful social experiences enhance the recall of previously acquired fear memories in male but not female mice. Activity-dependent tagging of cells in the dentate gyrus (DG) during fear learning revealed that these ensembles were endogenously reactivated during the social experiences in males. These reactivated cells were shown to be functional components of engrams, as optogenetic stimulation of the cells active during the social experience in previously fear conditioned animals was sufficient to drive fear-related behaviors. Our findings suggest that social encounters can reactivate pre-existing DG engrams and thereby strengthen discrete memories.

**Significance Statement:** Social environments can bolster and protect cognitive abilities. However, the relationship between social stimuli and individually learned memories remains enigmatic. Our work reveals that exposure to a stressed, naïve non-familiar conspecific or to the ambient olfactory-auditory cues of a recently stressed familiar conspecific induces reactivation of the cellular ensembles associated with a fear memory in the hippocampus. Artificially activating the hippocampal ensemble active during the social experience induces fearful behaviors only in animals that have previously acquired a negative memory, suggesting a fear-driving function of the reactivated ensembles and demonstrating the interaction between individual history and social experience. The neural resurgence of fear-driving ensembles during social experiences leads to a context-specific enhancement of fear recall. Our findings provide evidence that unlike directly physical stressors, ambient social stimuli can reactivate and amplify an individual’s memories.

## Results and Discussion

### Post-learning social stress amplifies behavioral expression of fear recall in a sex-dependent manner

To assess how social experiences influence existing memories, male and female mice were subjected to social or non-social salient events in the day between contextual fear conditioning (FC) and memory recall tests (**Fig. 1A**). A socially salient experience was provided in various forms: interaction with an unfamiliar juvenile intruder placed in the subject’s homecage (25, 26) to create a salient and mildly stressful event (Juvenile Intruder group); interaction with a recently shocked cagemate in the homecage to simulate a stressed familiar conspecific (Full Interaction group); indirect non-visual and non-physical exposure to a recently shocked cagemate in the homecage behind a 1-way mirror that permits unidirectional visual access for the shocked mouse to see the cagemates on the other side to simulate ambient exposure to conspecifics’ stress (1-Way Mirror group, **Fig. S1**); exposure to a recently shocked cagemate in the homecage through the same mirror covered in opaque black material to block both sides’ visual access (Opaque group). A non-social, salient experience was provided in the form of individual restraint stress in a tube (Restraint group), while a social but non-stressful experience was provided via interaction with a cagemate in the homecage (Cagemate control) or with a female mouse (Female Exposure) (**S Methods**). Additional control groups included mice that did not experience any events in the day between FC and recall (Neutral group), and mice that were exposed solely to the novel objects presented in the experimental groups without the associated salient events: the 1-way mirror object (Mirror No Mouse group) and the restraint tubes (Tubes group) to determine the impact of the novelty component (**S Methods**).

**Figure 1.**
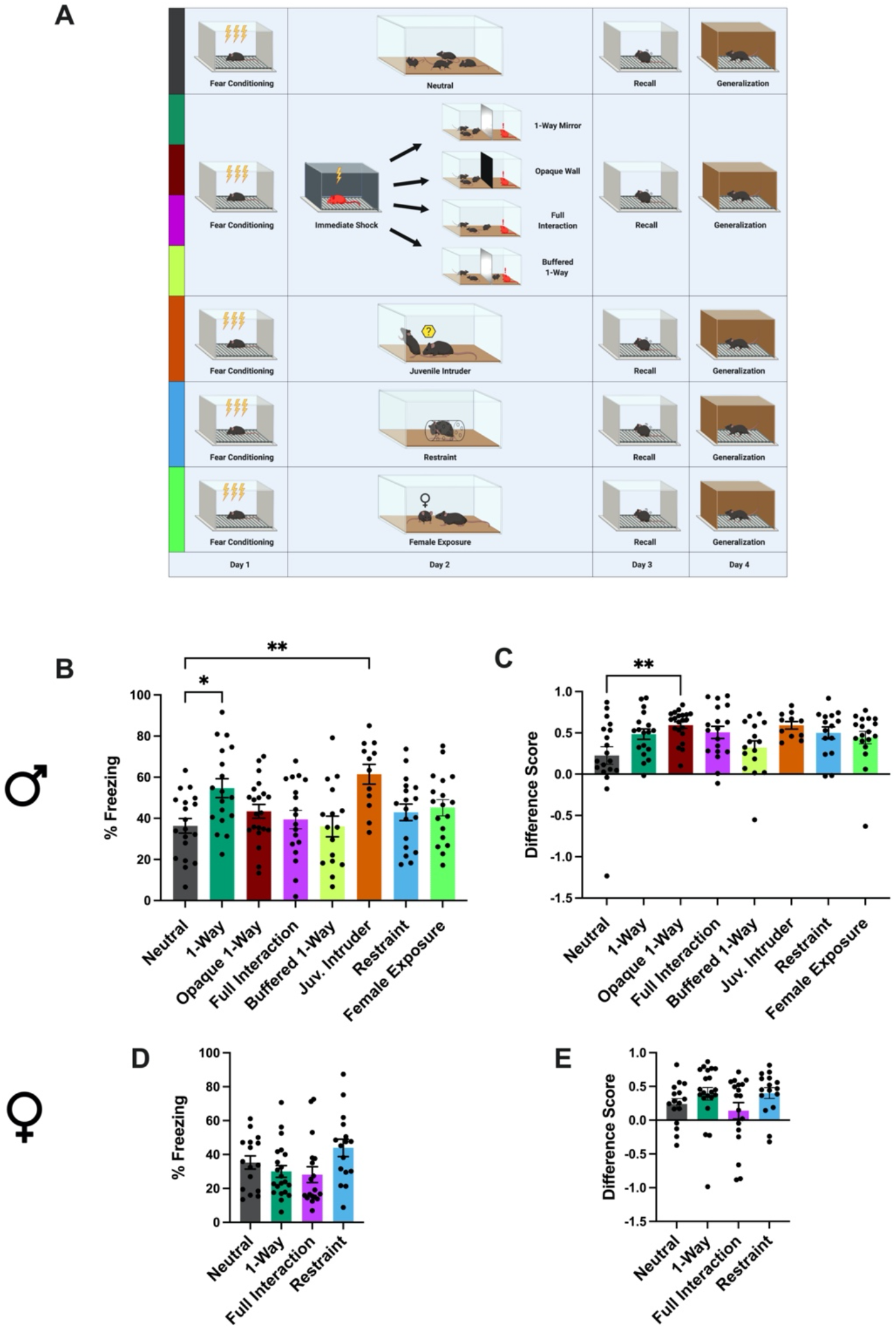
Social stimuli modulate subsequent fear recall in a sex-dependent manner. **A**) Schematic representation of behavioral schedule. **B** and **D**) Freezing levels during a 5 minute fear recall test for males (**B**) and females (**D**). Males: 1-Way ANOVA*F*_(7, 132)_=3.841, p=0.0008; Holm-Sidak’s multiple comparisons: ***p=0.0011 for Neutral vs Juvenile Intruder; **p=0.0114 for Neutral vs 1-Way Mirror. Females: 1-Way ANOVA*F*_(3, 67)_=2.626, p=0.0575. **C** and **E**) Differences scores, defined as [(freezing in recall test)-(freezing in generalization test)]/(freezing in recall test) for males (**C**) and females (**D**). Males: Kruskal-Wallis statistic=17.40, *p=0.0150; Dunn’s multiple comparisons: **p=0.0041 for Neutral vs Opaque. Females: ANOVA*F*_(3, 67)_=1.691, p=0.1773). Data shown as mean +/− SEM. See also **Fig S2** and **S3**.

Males exhibited higher levels of baseline freezing and froze more than females after the first two shocks of fear conditioning, after which freezing levels were similar (**Fig. S2B**), corroborating tone-cued experiments that report higher levels of freezing in males during acquisition (27). For males, all control groups exhibited similar freezing during recall and generalization tests to the Neutral group (**Fig. S2C and D**), indicating that the novel objects, as well as the non-stressful social experiences, did not impact recall. The Restraint group also performed similarly during recall and generalization tests to the Neutral group, suggesting that direct physical stress does not alter a previously established fear memory. We chose a short bout of immobilization (2 minutes) to match the intensity of stress with the much milder socially stressful experiences. However, a longer bout of immobilization (30 minutes) has previously been shown not to impact fear recall if applied 90 minutes post fear conditioning(28). Both the Juvenile Intruder and 1-Way Mirror groups froze more than the Neutral group during the recall test (**Fig. 1B**), despite similar levels of generalization (**Fig. 1C**), indicating that specifically social events enhance recall. These results extend the recent finding that observing a conspecific rat or human undergo an unconditioned stimulus (US) of electric shock reinstates context-specific fear memory^28^, suggesting that a conspecifics’ non US-specific stressed state can also strengthen observers’ fear memory.

We found that there was a subsequent enhancement of recall only if the stressed cagemate could see the experimental mice through a 1-way mirror; the Opaque group (in which a wall obscured the stressed cagemate’s visual access) froze the same as the Neutral group during recall, but exhibited higher difference scores, suggesting reduced generalization (**Fig. 1C**). The blocking of enhanced recall when the stressed cagemate can not see the other mice suggests the intriguing possibility that distressed mice emit different auditory-olfactory stimuli based on their perceived social context; i.e., producing more distressed signals when familiar conspecifics are nearby and available to buffer this distress(29).

The Full Interaction group also froze the same as the Neutral group during recall, indicating that physical interaction between the experimental mice and stressed cagemate blocked the effect on recall. It is crucial to note that physical interaction with a stressed familiar cagemate is not simply a heightened version of either the passive exposure to stressful emissions or of the interaction with an unfamiliar juvenile intruder mouse. In light of recent work on social buffering (30, 31) especially via physical interactions such as allogrooming(29), we hypothesized that full interaction permitted the experimental mice to buffer the shocked cagemate’s distress, and the subsequently reduced stress response failed to evoke the strengthening of cagemates’ fear recall. To test this hypothesis directly, we repeated the 1-Way Mirror protocol but placed one of the cagemates on the same side as the shocked mouse, thus allowing social buffering of the shocked mouse but maintaining the same conditions for the experimental mice on the other side (Socially Buffered group). As predicted, the Socially Buffered group froze the same as the Neutral group, indicating that the social buffering permitted in the Full Interaction group is sufficient to explain the blocking of effect on the experimental mice’s recall.

Females, on the other hand, exhibited no effect of either social or non-social experience on freezing during recall (**Fig. 1D**). The only difference in memory occurred for generalization of fear; both Mirror No Mouse and Tube Control groups exhibited elevated difference scores compared to the Neutral group (**Fig. S2F**), indicating that when presented in the absence of other salient experience, novel object exposure following fear conditioning can reduce the generalization of fear in females. The lack of effect of social stressors on females could be due to either a difference in cues emitted by the shocked females compared to the shocked males, to a difference in the response of cagemates to these cues, or a combination of both. Further experiments analyzing the ultrasonic vocalizations and chemical cues would enable clarification of this sex difference and testing of our hypothesis that male stressed mice emit different cues when unable to perceive social context behind the Opaque Wall.

These results demonstrate that intervening experiences between fear conditioning and recall can impact the intensity and context-specificity of fear expression in a sex-dependent manner. Social exposure to conspecifics’ distress, but not direct physical stressors, can enhance fear recall in males but not females. It is important to note that in group-housed animals, a memory acquired via fear conditioning could incorporate social stimuli produced in the homecage following conditioning. These social cues associated with the fear memory can feasibly act as a reminder cue, facilitating the enhancement of individually acquired fear memory via conspecifics’ stress. Although it is possible to exclude the putative social cues associated with a fear memory by singly housing mice after fear conditioning, this would also cause several confounds: exposure to intruders or shocked mice could be a qualitatively different experience for socially isolated mice, and social isolation induces a depressive phenotype that alters affective processing. Conceptually, if these social cues are necessary for the observed effect of social stress on fear memory, fear memories formed in group-living animals would similarly incorporate surrounding social cues and thus be similarly vulnerable to future enhancement via social stressors.

### Socially salient events reactivate ensembles previously active during fear conditioning

To test the hypothesis that fear memories are reactivated in the DG during the social experience for males and not for females, we used a cFos-based activity-dependent viral strategy (**Fig. 2A**) to track fear ensembles processed in the DG (dorsal DG; see methods) and BLA (the basolateral nucleus proper; see methods) post-fear conditioning (representative images, **Fig. 3B, Fig. S4**). As predicted, females did not exhibit reactivation of the DG fear trace during the social experience (**Fig. 2F**), while males exhibited significant reactivation during both types of social experience (for 1-Way Mirror 4.8x greater than chance, for Juvenile Intruder 4.3x greater than chance) (**Fig. 2D**). For males, Juvenile Intruder groups had more active cells, as identified by cFos expression, than 1-Way Mirror groups in both the DG and BLA (**Fig. 2D**). The increased level of DG activity during exposure to a juvenile intruder compared to a stressed cagemate behind a 1-way mirror, (**Fig. 3D**) despite both experiences occurring in the animal’s homecage, highlights the DG’s role in encoding experiences beyond physical contexts per se (32). We posit that the higher level of activity in the BLA and DG during exposure to a juvenile intruder reflects the richness of direct social interaction during the juvenile intruder’s presence that is not permitted in the 1-way mirror paradigm.

**Figure 2.**
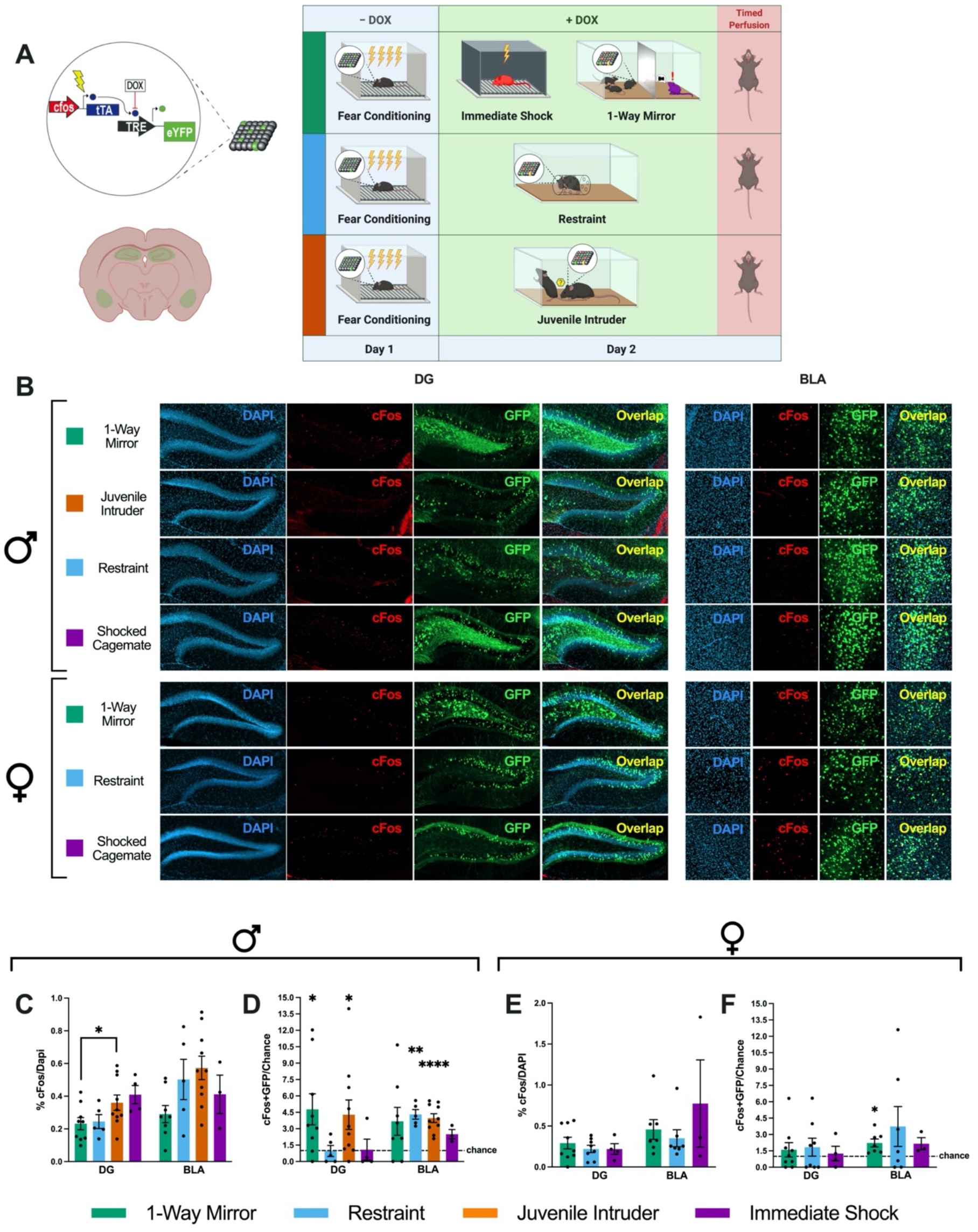
Socially salient but not directly stressful experiences reactivate pre-existing fear ensembles. **A)** Schematic representation of the doxycycline mediated viral tagging construct (left) and experimental design (right). **B**) Representative confocal images of DG and BLA histology visualizing cFos-tTa+Tre-EYFP positive cells (active during FC, green) and cFos positive cells (active during the different types of stress exposure, red). **C** and **E**) Percentage of cFos positive cells over DAPI positive cells across brain regions in males and females (Males: DG - 1Way ANOVA*F*_(3, 24)_=3.028, p=0.0490, Holm-Sidak’s multiple comparisons: *p=0.0339 for Juvenile Intruder vs 1-Way Mirror; BLA - 1Way ANOVA*F*_(3, 22)_=2.710, p=0.0697. Females: DG - 1Way ANOVA*F*_(2, 18)_=0.4501, p=0.6445; BLA - ANOVA*F*_(2,14)_=c0.9580, p=0.4074). **B** and **D**) Degree of overlap between the two labeled populations normalized over chance (%cFos/DAPI * %EYFP/DAPI). 1-sample t-tests. Males: DG - Juvenile Intruder t=2.436, df=9, *p=0.0376; 1-Way Mirror t=2.656, df=8, *p=0.0290. BLA - Juvenile Intruder t=7.411, df=9, ****p<0.0001; Restraint t=7.462, df=4, 0.0017, **p=0.0017. Females: BLA - 1-Way Mirror t=3.387, df=6, *p=0.0147). See details for non-significant results in supplementary text. Data shown as mean +/− SEM. Dotted line represents chance. See also **Fig S4**.

**Figure 3.**
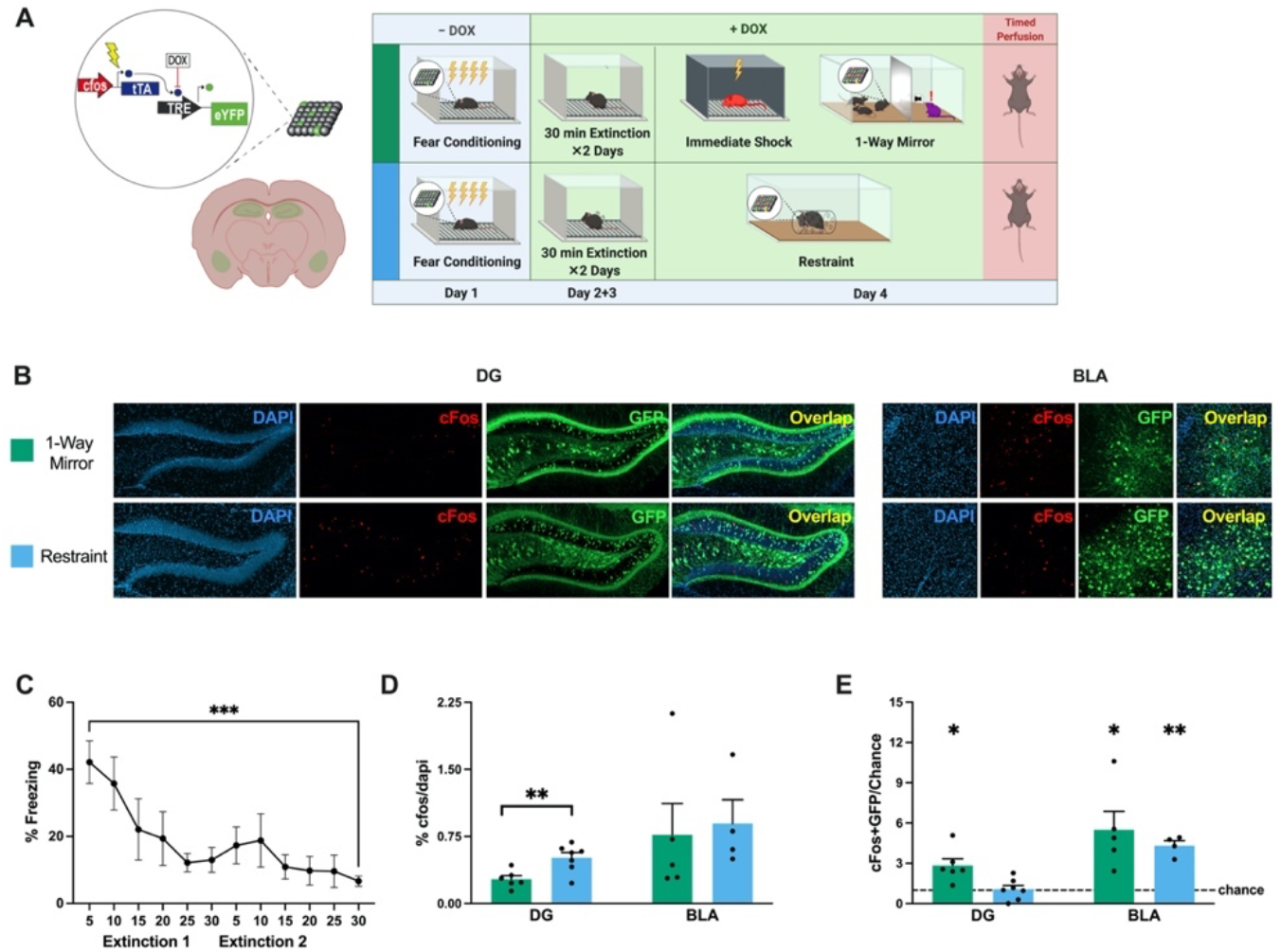
Socially salient experiences reactivate fear ensembles post fear extinction. **A)** Schematic representation of the doxycycline mediated viral tagging construct (left) and experimental design (right). **B)** Confocal visualization of DG and BLA histology showing cFos-tTa+Tre-EYFP labelled cells (active during FC, green) and cFos+ cells (active during the different types of stress exposure, red). **C)** Freezing throughout extinction training (RM 1Way ANOVA*F*_(3.230, 25.84)_=5.165, p=0.0054, Holm-Sidak’s multiple comparisons for first vs last 5 minutes, n=9, t=5.688, ***p=0.0005). **D)** Percentage of cFos+ cells per DAPI+ cells (DG - t-test, t=3.268, df=11, **p=0.0075; BLA - t-test, t=0.2716, df=7, p=0.7938). **E)** Degree of overlap between the two labeled populations normalized over chance (%cFos/DAPI * %EYFP/DAPI). 1-sample t-tests. DG - Restraint t=0.195, df=6, p=0.8516; 1-Way Mirror t=3.67, df=5, *p=0.0153. BLA - Restraint t=9.031, df=3, **p=0.0029; 1-Way Mirror t=3.257, df=4, *p=0.0312). Dashed line represents chance. Data shown as mean +/− SEM.

Neither sex demonstrated DG fear trace reactivation during the non-social stressors of restraint or immediate shock (**Fig. 2D, left**), and previous work shows that fear trace reactivation does not occur in neutral contexts (33), supporting the socially-specific nature of this reactivation. Conversely, in the BLA, both Juvenile Intruder and Restraint groups displayed reactivation of tagged cells in males (**Fig. 2D, right**), suggesting that the common denominator of direct physical stress in the form of restraint or interaction with an intruder might recruit amygdalar ensembles also activated during fear conditioning the previous day. However, females in the 1-Way Mirror group also displayed a modest but significant reactivation of tagged cells in the BLA (i.e. ~2x greater than chance, whereas the above-mentioned males displayed ~4x greater than chance overlap). Although our immediate-early gene mediated resolution of the reactivated population does not permit within-region subdivision of functionally distinct subsets of a fear engram, the result that the BLA ensemble is more active than the DG ensemble during other experiences suggests that these reactivated components of the BLA fear ensemble are involved in processes other than driving acute fear responses. This idea is further supported by the discorrelation of overlaps in the BLA from overlaps in the DG in both sexes (Supplementary Text) and corroborated by the optogenetic stimulation experiment described below (**Fig. 4)**.

**Figure 4.**
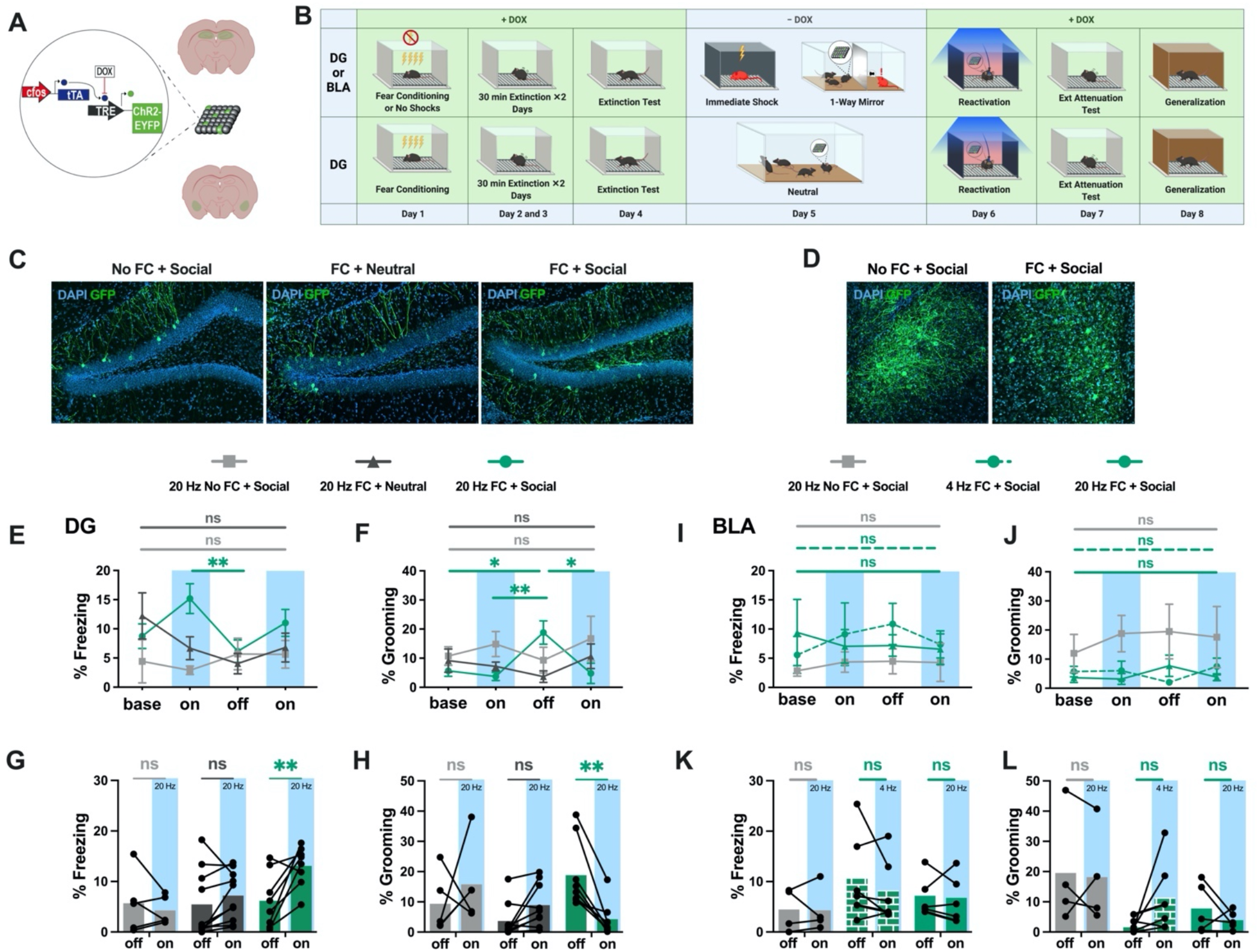
DG ensembles active during social stress drive fear only in previously fear conditioned mice. **A)** Schematic representation of activity-dependent engram tagging strategy. **B)** Behavioral schedule. **C** and **D)** Representative images of DAPI+ cells (blue) and Chr2-EYFP+ cells (green) in the DG (**C**) and BLA (**D**). % freezing levels throughout optogenetic stimulation of cells active in DG (**E**) and BLA (**I**): 2-Way ANOVA, Holm-Sidak’s multiple comparisons: for FC’d DG first light-on vs light-off t=3.821, *p=0.0385. % freezing levels for the light-off and the averaged light-on sessions in DG (**G**) and BLA (**K**): 2-Way ANOVA, Holm-Sidak’s multiple comparisons: for FC’d DG light-on vs light-off t=4.072, **p=0.0018. % self-grooming levels throughout optogenetic stimulation in DG (**F**) and BLA (**J**): 2-Way ANOVA, Holm-Sidak’s multiple comparisons: for FC’d DG for light-off vs baseline t=2.897, *p=0.0211; vs first light-on t=3.330, **p=0.0091; vs second light-on t=2.09, *p=0.0153. % self-grooming levels for the light-off and the averaged light-on sessions in DG (**H**) and BLA (**I**): 2-WAY ANOVA, Holm-Sidak’s multiple comparisons: for FC’d DG for light-on vs light-off t=3.5578, **p=0.006. Data shows as mean, in E, F, I, J +/− SEM. Detailed statistics shown in Supplemental Text. See also **Fig S5**.

To test whether the reactivation of the fear trace in males occurred only during events happening directly after fear conditioning, we also tracked reactivation of the fear memory following a two day extinction protocol (**Fig. 3A,** representative images in **3b**), after which mice froze significantly less in the fear context (**Fig. 3C**). The percentage of cFos+ cells in DG was slightly but consistently higher during non-social restraint stress than during the social experience (**Fig. 3D, left**), indicating a higher level of overall DG activity. Similar to pre-extinction, the DG fear ensemble was reactivated during the social experience but not during non-social restraint, and the BLA fear ensemble was reactivated during non-social restraint (**Fig. 3E**). Unlike our pre-extinction results, post-extinction the BLA fear ensemble was also reactivated during the 1-Way Mirror experience (**Fig. 3E**). It is important to note that our data indicate that a significant portion of the fear ensemble was reactivated and may not reflect whether the reactivated subset of cells are the same or different across these experiences.

Collectively these data suggest that specifically social situations reactivate subsets of fear ensembles in the DG for males and not females, and do so even after extinction training. Strikingly, the degree of DG cellular reactivation during the social encounter in males both before and after extinction was similar to previously reported results in animals re-exposed to the original fear context (34). These results are consistent with the theory that endogenous reactivation of fear ensembles enhances recall (35) and is the mechanism by which multiple exposures to the contexts (36) or cues (37) present during conditioning can strengthen memories in the days after learning. We show that this reactivation does not occur during non-social physical stress, and previous work has shown that it does not occur in neutral contexts (33), supporting the idea of a causal relationship between reactivation and enhanced recall. Given the malleability of engrams (38, 39), the natural reactivation of a fear memory ensemble during a stressful social experience could plausibly add additional negative valence from the current experience to the representation of the former, explaining why a single negative social encounter enhances fear whereas multiple exposures to associated neutral contexts (36) are necessary for enhancement.

### Stressful social situations activate fear-driving DG ensembles in previously fear conditioned mice

Next, we aimed to investigate the functional properties of the cells which were active during exposure to a stressed cagemate in previously fear-conditioned animals. Fear ensemble reactivation in the DG was correlated specifically with the social experiences that heightened recall, while reactivation in the BLA was more variable. Thus, we hypothesized that the fear ensembles reactivated in the DG were comprised of fear-activating cells while the ensembles reactivated in the BLA were comprised of non fear-driving components of the ensemble. We predicted that the fear ensembles reactivated during social stress in the DG, but not in the BLA, would cause the cells tagged during this period to drive fear behaviors when optogenetically stimulated. To this end, we tagged DG or BLA cells with channelrhodopsin-2 during our 1-way mirror paradigm or during a neutral home-cage period (**Fig. 4a-d;** See Methods). To ensure that we were not capturing a persistent fearful state but rather the re-emergence of a fear ensemble, we chose to first employ a two day extinction protocol before tagging the cells active during the social or neutral experience. As we show in Figure 3, this timing still results in reactivation of the fear conditioning ensemble. The following day, the tagged ensembles were stimulated in a novel context to test their capacity to drive fearful behavior.

In support of our hypothesis, 20Hz stimulation of the DG ensembles active during the social experience in previously fear conditioned animals drove higher levels of freezing during the light-on sessions (**Fig. 4J**) as well as increased self-grooming (**Fig. 4K**) in the light-off period following stimulation, which is thought to reflect a de-arousing behavioral state(40). To ensure that this effect was not due simply to negatively valenced cells active during social stress or active post-FC regardless of social experience, we showed that fear behaviors were not induced by either stimulation of cells active in a social setting without prior fear conditioning, or of cells active in a neutral setting after fear conditioning (**Fig. 4, 4K**). The size of the labeled ensembles did not differ between any of these three groups (**Fig. S3E**), indicating that DG activity levels were generally similar and it was not the number of tagged cells but their functionality as fear driving cells that induced fear. In the context of the social experience, the functional capacity of the active ensembles in the DG to drive obvious fear behaviors was quiescent (example **Movie S1** and **Movie S2**), however, we postulate that the more nuanced documented deficits in social interaction following trauma (41–44) could in part be driven by DG fear ensemble activation.

The coactivation of ensembles leads to co-allocation and the linking of memories, as shown in both mouse (45) and human (46) studies. Our data suggests that during salient social experiences, the DG is concurrently reactivating a pre-existing fear engram and processing the current social stimuli (**Fig. 2**). We therefore posit that the reactivated subset of the fear ensemble was co-allocated to form part of the ensemble encoding the social encounter, perhaps explaining why the total number of cells active during the social exposure was similar regardless of prior fear conditioning (**Fig. 4I**) and why stimulation drove fearful behaviors only in the previously fear conditioned animals (**Fig. 4J and 4K**).

On the other hand, the number of tagged cells in the BLA was higher during social stress if mice had been previously fear conditioned **(Fig. S3F**), suggesting that the combination of prior fear conditioning and social stress leads to heightened amygdalar activity. This is consistent with the notion that sustained amygdalar hyperactivity underlies persistent potentiation of fear and anxiety after stressful events (47); in rats, prior stress reduces inhibitory control in the BLA, increasing excitability and plasticity (48). Although stimulation of the BLA fear engram drives freezing(20), counterbalanced 20Hz and 4Hz stimulation of cells tagged during social stress did not cause freezing or grooming (**Fig. 4H**), indicating that the amygdalar cells that were active during both FC and social stress (**Fig. 3E**) did not comprise the portion of the original fear engram that drives freezing.

Collectively, these results indicate that the DG ensemble activated during exposure to a stressed cagemate includes a previously established functional fear engram, while the active BLA ensemble is not composed of fear-driving cells.

## Conclusion

Memory recall is a malleable phenomenon that can be altered by intervening experiences, as associated context or cues can reactivate engrams and thus make them labile during incubation (35–37, 49). In this study, we discovered that even in the absence of associated reminders, salient social encounters can strengthen a pre-existing fear memory in males, but not in females. We explored the role of endogenous cellular reactivation of fear conditioning ensembles in the hippocampus and amygdala during subsequent social vs non-social salient events. While the two forms of socially salient events induced reactivation of the fear ensemble in the DG, the two forms of non-social physical stress did not cause reactivation. Unlike in the DG, reactivation in the BLA was not associated with the social, memory-enhancing experiences. To determine the functional nature of the ensembles activated during the social experiences, we optogenetically tagged these cells and artificially stimulated them with light in neutral conditions. We found that light-induced stimulation caused fear behaviors only when mice had been previously fear conditioned and when the cells stimulated were tagged during the social experience, suggesting that the ensembles reactivated by socially salient events are fear-driving engrams. Thus, we propose that DG engram reactivation provides a mechanism for stressful social experience to potentiate negative memories. Future studies are necessary to confirm the causal role of engram reactivation with an activation-blocking strategy such as chemogenetics while avoiding potentially confounding compensatory effects(50), (51)).

For both humans and mice, healthy social conditions include not only positive but also moderately stressful interactions. While extreme social stress can impair memory capabilities (8), a rich social life can protect (7) and enhance (5) memory. Although this enriching effect on learning and memory is often discussed as a function of positive social support, our findings suggest that mildly negative social exposure to conspecifics’ stress also plays an important role in strengthening pre-existing memory. From an evolutionary perspective, a stressed conspecific or their ambient auditory-olfactory emissions might signal a dangerous situation in which it is adaptive to hone one’s own memories to guide decision making. Altogether, our findings provide evidence that the mnemonic contents of a social animal’s brain are modulated throughout the course of social experiences, and we anticipate ensuing studies to unravel the circuitry bridging social sensory inputs and pre-existing hippocampal engrams.

## Supporting information

1 way mirror

Resident Intruder

## Acknowledgments

Behavioral timelines created with Biorender.com. The authors would like to thank Dr. Susumu Tonegawa and his lab for providing the activity-dependent virus cocktail. **Funding:** This work was supported by an NIH Early Independence Award (DP5 OD023106-01), an NIH Transformative R01 Award, a Young Investigator Grant from the Brain and Behavior Research Foundation, a Ludwig Family Foundation grant, the McKnight Foundation Memory and Cognitive Disorders Award, and the Center for Systems Neuroscience and Neurophotonics Center at Boston University. **Author contributions:** Conceptualization: ABF, HL, RC, and SR; investigation: ABF, HL, RC, TG, AV; methodology: ABF, HL, TG, YZ; resources: SR; supervision: ABF, SR; visualization: ABF, HL, RC; formal analysis: ABF and HL; writing - original draft: ABF, RC, HL; writing - review and editing: all authors; and funding acquisition: SR. **Competing interests:** Authors declare no competing interests; and **Data and materials availability:** All data is available in the main text or the supplementary materials.

## Materials and Methods

### Surgery

For all surgeries, mice were initially anesthetized under 3.5% isoflurane inhalation and then maintained during surgery at 1.0-2.0% isoflurane inhalation through stereotaxic nosecone delivery. Ophthalmic ointment was applied to both eyes to provide adequate lubrication and prevent corneal damage. The hair above the surgical site was removed with scissors and subsequently cleaned with alternating applications of betadine solution and 70% ethanol. 2.0% Lidocaine HCl was injected subcutaneously as local analgesia prior to 10-15mm midsagittal incision of the skin. For optogenetic implant surgeries, two bone anchor screws were secured into the cranium, one anterior and one posterior to the target injection and fiber placement sites. All animals then received bilateral craniotomies with a 0.6 mm drill-bit for dorsal dentate gyrus (dDG) and basolateral amygdala (BLA) injections. For all dDG and BLA surgeries, a 10μL air-tight Hamilton syringe with attached 33-gage beveled needle was lowered to the coordinates for DG of −2.2mm anteroposterior (AP), ±1.3mm mediolateral (ML), −2.0mm dorsoventral (DV), and for BLA of −1.35mm (AP), ±3.25mm mediolateral (ML), and −5.0mm (DV). All coordinates are given relative to bregma (mm). For overlap surgeries, a volume of 300 nL of AAV9-cFos-tTa + TRE-EYFP viral cocktail was bilaterally injected at 200 nL/min using a micro-infusion pump for each coordinate (2×300 nL for dDG and 2×300 nL for BLA) (UMP3; World Precision Instruments). After injection was complete, the needle was left in place 2 minutes prior to incremental retraction of the needle from the brain. For dDG **or** BLA optogenetic surgeries, a 300nL viral cocktail of AAV9-cFos-tTa + AAV9-TRE-ChR2-EYFP was bilaterally injected into the dDG **or** BLA (separate surgeries and animals for each brain region; see overlap surgeries for coordinates and procedure for injection). Following viral injection, bilateral optical fibers (200μm core diameter; Doric Lenses) were placed 0.4mm above the injection sites (dDG: −1.6mm DV; BLA: −4.6mm DV). The implants were secured to the skull with a layer of adhesive cement (C&M Metabond) followed by multiple layers of dental cement (Stoelting). Mice were injected with a 0.1 mg/kg intraperitoneal dose of buprenorphine (volume dependent on weight of animal) and placed in a recovery cage atop a heating pad until recovered from anesthesia. Viral targeting was confirmed by histological study and only animals with proper viral expression were utilized for data analysis.

### Immunohistochemistry

Mice were overdosed with 3% isoflurane and perfused transcardially with cold (4°C) phosphate-buffered saline (PBS) followed by 4% paraformaldehyde (PFA) in PBS. Brains were extracted and kept in PFA at 4°C for 24-48 hours and then transferred to PBS solution. Brains were sectioned into 50μm thick coronal sections with a vibratome and collected in cold PBS. Sections were blocked for 1-2 hours at room temperature in PBS combined with 0.2% triton (PBST) and 5% normal goat serum (NGS) on a shaker. Sections were incubated in primary antibody (1:1000 rabbit anti-c-Fos [SySy]; 1:5000 chicken anti-GFP [Invitrogen]) made in PBST-NGS at 4°C for 48 hours. Sections then underwent three washes in PBST for 10 minutes each, followed by a 2 hour incubation period at room temperature with secondary antibody (1:200 Alexa 555 anti-rabbit [Invitrogen]; 1:200 Alexa 488 anti-chicken [Invitrogen]) made in PBST-NGS. Sections then underwent three more wash cycles in PBST. Sections were mounted onto microscope slides (VWR International, LLC). Vectashield HardSet Mounting Medium with DAPI (Vector Laboratories, Inc) was applied and slides were coverslipped and allowed to dry overnight. Once dry, slides were sealed with clear nail polish on each edge and stored in a slide box in the fridge. If not mounted immediately, sections were stored in PBS at 4°C.

### Behavior

#### Fear Conditioning

Fear conditioning (FC) for all experiments took place in a 18.5 × 18 × 21.5 cm chamber with aluminum side walls and plexiglass front and rear walls (Context A). Each cage animal was placed in a separate chamber (4) and received shocks in parallel during the session. The session consisted of 4 shocks over a time span of 500s (shocks at 198s, 278s, 358s, 438s). Depending on the experiment, the length and strength of the shock varied. For the experiments without extinction training (recall 24hrs after social session), the four shocks were 1 second each at 1mA. For the experiments with extinction, in order to obtain homogenous and reliable fear responses at a more remote time point, more intense fear conditioning was utilized, with shocks at 1.5mA for 2 seconds. After the session, animals were placed back into their home cage and in an isolated holding area until all animals in a cohort had been fear conditioned.

#### Extinction Training

Animals were placed into Context A (see above) for a duration of 30 minutes on the day following fear conditioning. A second extinction training session occurred on the next day (30 minutes). All animals in a cage of four underwent extinction training simultaneously. After the session, animals were placed back into their home cage and in an isolated holding area until all animals in a cohort had undergone extinction training.

#### Extinction Test

For the extinction test, animals were placed into Context A for a duration of 5 minutes. Depending on the experiment, the five minute extinction test was used to assess successful extinction and occured on the day following the second session of extinction training. As in fear conditioning and extinction training, all animals in a cage of four underwent testing simultaneously and were placed back into their home cage and in an isolated holding area until all animals in a cohort were tested.

#### Recall Test

For the recall test, animals were placed into Context A for a duration of 5 minutes. As in fear conditioning and extinction training, all animals in a cage of four underwent testing simultaneously and were placed back into their home cage and in an isolated holding area until all animals in a cohort were tested.

#### Generalization

To assess generalization, 24 hours after the recall session animals were placed into Context B (see above for description) for a duration of 5 minutes. All animals in a cage of four underwent testing simultaneously and were placed back into their home cage and in an isolated holding area until all animals in a cohort were tested.

#### Juvenile Intruder/Cagemate Control

Animals were removed from the home cage and placed into a separate, clean cage with access to food and water. The lid of the cage and the feeder were removed from the home cage, which served as the interaction chamber. An experimental mouse was placed back into the home cage and allowed 1 minute to acclimate. An unfamiliar younger (p43-49) intruder mouse was placed into the home cage for ten minutes of interaction with the experimental resident mouse; younger mice were used to avoid confounding effects of mutual aggression (25, 52). A clear acrylic top was placed over the home cage during interaction. This was repeated for each experimental cage animal. A different intruder mouse was used in each session. After each session, both mice were removed from the home cage and placed into a separate holding cage for their respective group. For cagemate control experiments, a cagemate of the resident mouse was used for interaction in lieu of the unfamiliar intruder. The social session was conducted with a procedure otherwise identical to that used for RIT.

#### Immediate Shock

Immediate shock was administered in a 18.5 × 18 × 21.5 cm chamber with vertical black and white striped (3 cm width) sides, a plastic container holding gauze soaked in almond extract under the chamber floor, and red light as room illumination (Context B). The animal received a single 1.5mA shock if after extinction or 1mA shock if without extinction, beginning 1s after trial initiation and lasting 2s, followed by 57s in the chamber, for a total duration of 60s.

#### Full Interaction

The lid of the cage and the feeder were removed for social interaction and a clear acrylic top was placed over the home cage. The recently shocked cagemate was then placed into the cage and all animals were left in the testing room for one hour of social interaction. After one hour, the home cage was returned to its normal condition.

#### 1-Way Mirror/Mirror No Mouse Control/Opaque Control

Mirror inserts were created out of laminated cardboard and one way mirror material (OuBay). Wooden dowels were placed on the bottom to support the structure. Dimensions were based to fit tightly into the cage (7” x 11” x 5”) to separate the container into two sections (a smaller area for the recently shocked cagemate and larger area for the rest of the cagemates). The lid of the cage and the feeder were removed for social interaction and a clear acrylic top was placed over the home cage. In order to enhance the effect of the mirror by creating a large light difference, black covers were created to cover the section of the cage with the recently shocked cagemate. Refer to **Figure S1** for a visual representation of the set-up. After Immediate Shock, the recently shocked cagemate was immediately placed into the section opposite of and separate from the other cagemates. Animals were left in the testing room for one hour of social interaction. After one hour, the recently shocked cagemate was removed and immediately euthanized by overdose with sodium pentobarbital to ensure any subsequent effects were due to the manipulation rather than any social interaction afterward, and the home cage was returned to its normal condition. For the Mirror No Mouse Control, all procedural steps were identical to the 1-Way Mirror experiments, except that there was no shocked cagemate placed on the other side of the mirror. For Opaque control experiments, the 1-Way Mirror insert was altered through the addition of a black insert on the side of the shocked cagemate so that the wall was bidirectionally blocking visual input from the other side. Social interaction was conducted with a procedure otherwise identical to that used for 1-way mirror experiments.

#### Social Buffering

To determine the impact of social buffering of the shocked cagemate, subjects experienced the same protocol as 1-Way Mirror except that one of the cagemates was placed behind the mirror insert during the immediate shock, so that the shocked mouse was able to directly interact with this cagemate during the following 1 hour. Afterward, both the shocked cagemate and “bufferer” mouse were placed into a new cage before the insert was removed to ensure the effect on experimental mice were limited to the time window of the manipulation.

#### Restraint/Tube Control

One at a time, animals were removed from the home cage and enclosed in a plastic restraint tube containing holes to permit air flow. The tube was then placed in the center of a clean cage for a duration of two minutes. At the end of the two minutes, the animal was released from the tube into one separate clean cage where all the mice were reunited post restraint. For the tube control test, all animals remained in the home cage and a clean restraint tube was placed into the home cage for a duration of two minutes.

#### Female Exposure

One at a time, animals were removed from the home cage and placed into a separate, clean cage for 10 minutes of interaction with an unfamiliar female conspecific. To lessen the females’ stress induced resistance to mating, some of the females’ bedding was sprinkled into the cage and the females were habituated to the new environment for 1 minute prior to introduction of the experimental male. A clear acrylic top was placed over the cage during interaction. This was repeated for each experimental cage animal. A different female was used in each session. After each session, both mice were removed from the interaction cage and placed in a new cage, so that naive animals could not interact with those that had already gone through the experience.

### Optogenetic Stimulation

The optogenetic stimulation session occurred 24 hours after the tagging experience (1-Way Mirror). Prior to the start of the session, optogenetic patch cords were tested to ensure a minimum 10mA laser output. Mice were given stimulation in a separate room than Context A or B. For stimulation, the mice were attached to the optogenetic cords in the palm of ABF’s hand and placed into a striped acrylic chamber with either white or dimmed white + red light and either almond or orange scent (Context C). The session lasted 8 minutes and consisted of 4 alternating 2 minute periods of laser stimulation (off/on/off/on; dDG: 450 nm, 20 Hz, 10 ms pulse width(53); BLA: 450 nm, 4 Hz and 20 Hz, 10 ms pulse width(54)). The BLA group received 4 Hz and 20 Hz stimulation counterbalanced in different contexts (orange vs almond scent, white vs red light) separated by about 1.5hrs.

### Behavior Scoring

#### FreezeFrame

Videos of behavioral sessions were obtained using cameras secured to the chamber either above or to the side of the subject. For extinction, extinction test, and recall sessions, FreezeFrame/View (Coulbourn Instruments; Whitehall, PA) was used to score freezing behavior, defined as 1.25 seconds of animal immobility.

#### ezTrack

Videos of behavioral sessions were obtained using cameras secured to the chamber either above or to the side of the subject. Videos of the open field test were obtained using a webcam secured above for an aerial view. ezTrack(55) was used to analyze all open field videos. An ROI of the center was defined manually according to the lines present on the floor of the open field arena. The software tracked the movement of the mouse and calculated the amount of frames in which the mouse moved less than specified parameters (both pixel change and distance traveled). The software then tracked and calculated the amount of frames the mouse spent in the center, which was converted to the amount of time spent in the center according to video parameters.

#### Manual scoring

For optogenetic sessions, due to cord movement and lighting conditions interfering with automated scoring, all freezing and grooming quantification was done manually. This was then converted to a percentage of time that the mouse spent freezing within the bins of stimulation in the session. Each video was scored for grooming by HL, and for freezing by 2 separate observers (ABF and HL or ABF and RC) whose scores were averaged to mitigate variation in bout length perception. Grooming was defined as any syntactic-chain cephalocaudal grooming, scratching, licking, or other observable forms of non-chain self-grooming performed by the animal(40). Freezing was defined as any observable complete cessation of movement, other than breathing, by the animal.

### Imaging and Cell Counting

All coronal brain slices were imaged through a Zeiss LSM 800 epifluorescence microscope with a 20x/0.8NA objective using the Zen2.3 software. Images of the basolateral amygdala were captured in a 2×2 tile (640×640 microns) Z-stack. Images of the dentate gyrus were captured in a 4×2 tile (1280×640 microns) Z-stack. DAPI and GFP were imaged as separate channels for target verification and ensemble size quantification. For overlap counts, DAPI, cFos, and GFP were imaged (DAPI and cFos simultaneously and GFP as a separate channel). 3-4 different slices were imaged for each animal for averaging.

#### FIJI Software

Images were processed for greater clarity before quantification. The Subtract Background tool was used to enhance contrast of cells to background, and the Despeckle was used to minimize noise that may interfere with quantification. ROIs were selected using Freehand Selection so that only cells within the brain region would be analyzed. The DAPI channel was segmented using a custom pipeline involving 3D Object Counter. The cFos and GFP channels were initially segmented using a custom pipeline utilizing the 3D Iterative Thresholding tool of 3D ImageSuite (56). Once cells segmented, the z-stacks were Z-Projected into a single slice image and saved.

#### Cell Profiler

Segmented images of DAPI, cFos, and GFP were loaded into CellProfiler and run through a pipeline that identified cells with more stringent parameters of size and shape. For overlap quantification, the final step in the pipeline counted all identified objects between the cFos and GFP images that had an overlap of greater than 80%.

#### Cell Quantification and Analysis

Once cells were counted, the relative amounts of cFos+ only, GFP+ only, and cFos+GFP+ (overlap) cells in each slice was normalized over the total amount of DAPI cells present (cFos+/DAPI, GFP+/DAPI, Overlap/DAPI). Chance of an overlap was defined as (cFos+/DAPI)x (GFP+/DAPI). Overlap over Chance was then calculated by dividing Overlap/DAPI by the Chance Overlap calculated in the previous step.

### Data Analysis

#### Freezing during recall and Generalization

Behavioral and histological data were analyzed using GraphPad Prism version 9.1.1 for Mac OS, (GraphPad Software, San Diego, California USA). For the recall behavior, we aimed to compare freezing levels between a neutral control group and each manipulated group. As standard deviations passed Brown-Forsythe’s and Bartlett’s tests for non-significantly different standard deviations and all normality tests, we conducted ordinary one-way ANOVAs followed by post-hoc comparisons to the Neutral group using the Holm-Sidak’s correction for multiple comparisons. For generalization behavior, we compared the generalization scores ((recall freezing - contextB freezing)/recall freezing) of each group to those of the Neutral group. When distributions did not pass the Anderson-Darling test for normality, we conducted the Kruskal-Wallis test followed by Dunn’s multiple comparisons to the Neutral group.

#### Freezing and self-grooming during optogenetic stimulation

For the DG optogenetic data, we aimed to determine the effects and interactions of optogenetic stimulation, prior fear conditioning, and social experience during the tagging window. We therefore used two-way Repeated Measures ANOVAs followed by pair-wise multiple comparisons between stimulation trials within each of the three groups, with Holm-Sidak’s correction. These same tests were used for the BLA optogenetic data, where we aimed to determine the effects and interactions of optogenetic stimulation, frequency of stimulation, and prior fear conditioning.

#### Cellular overlaps

To quantify the degree of overlap in cellular populations active during fear conditioning and each of our conditions the following day, we aimed to compare the number of overlaps in each brain region to that expected by chance based on the size of the two populations. We therefore normalized overlaps over chance as described above in *Cell Quantification and Analysis* and compared to chance with 1-sample t-tests. In order to compare the levels of activity during each of the salient and control conditions, we also performed 1-way ANOVAs on % cfos/DAPI in each brain region, followed by Holm-Sidak’s multiple comparisons.

## Supplementary Text

### Statistical Analysis Details

**Figure 2**: Socially salient but not directly stressful experience reactivate pre-existing fear ensembles

**Fig. 2d**: Males, 1-sample t-test: DG Restraint t=0.0667, df=4, p=0.95; Shocked Cagemate t=0.085, df=3, p=0.937. BLA 1-Way Mirror t=2.101, df=7, p=0.0737; Shocked Cagemate t=3.492, df=2, p=0.0731.

**Fig. 2e**: Females, 1-Way ANOVA: DG *F*_2, 18_=0.4501, p=0.6445. BLA *F*_2,14_=0.9580, p=0.4074.

**Fig. 2f**: 1-sample t-test for females, DG Restraint t=1.022, df=7, p=0.3406; 1-Way Mirror t=0.9085, df=8, p=0.3902; Shocked Cagemate t=0.3829, df=3, p=0.7273. BLA Restraint t=1.496, df=6, p=0.1852; Shocked Cagemate t=2.084, df=2, p=0.1725.

**Correlations of DG to BLA reactivation,** Pearson Correlations: Males, 1-Way: n=8, Pearson r = 0.116, p=0.784; Restraint: n=4, Pearson r = 0.869, p=0.13; Juvenile Intruder: n=9, Pearson r=0.530, p=0.142. Females, 1-Way: n=6, Pearson r=-0.266, p=0.610; Restraint: n=5, Pearson r=-0.139, p=0.823.

**Figure 4**: DG ensembles active during social stress drive fear only in previously fear conditioned mice

**Fig. 4e**: 2-Way RM ANOVA light x group *F*_6, 57_=1.628, p=0.1560; light *F*_1.917, 36.43_=0.9090, p=0.4081; group *F*_62, 19_=3.959, p=0.0365.

**Fig. 4f**: 2-Way RM ANOVA light x group *F*_6, 57_=3.071, p=0.0113; light *F*_3, 57_=0.3763, p=0.7704; group *F*_2, 19_=1.645, p=0.2193.

**Fig. 4g**: 2-Way RM ANOVA light x group *F*_2, 20_=5.120, p=0.0160; group *F*_2, 20_=1.801, p=0.1909; light *F*_1, 20_=5.315, p=0.0320.

**Fig. 4h**: 2-Way RM ANOVA light x group *F*_2, 19_=7.889, p=0.0032; group *F*_2, 19_=2.007, p=0.1619; light *F*_1, 19_=0.1406, p=0.7118.

**Fig. 4i**: 2-Way RM ANOVA light x group *F*_6, 36_=0.3690, p=0.8938; light *F*_3, 36_=0.1871, p=0.9045; group *F*_2, 12_=0.7903, p=0.4760.

**Fig. 4j**: 2-Way RM ANOVA light x group *F*_6, 36_=1.266, p=0.2971; light *F*_2.559, 30.71_=0.6596, p=0.5602; group *F*_2, 12_=3.119, p=0.0812.

**Fig. 4k**: 2-Way RM ANOVA light x group *F*_2, 12_=0.5180, p=0.6084; group *F*_2, 12_=1.062, p=0.3762; light *F*_1, 12_=0.8643, p=0.3709.

**Fig. 4l**: 2-Way RM ANOVA light x group *F*_2, 12_=2.285, p=0.1443; group *F*_2, 12_=2.503, p=0.1235; light *F*_1, 12_=0.2462, p=0.6287.

---

**Fig. S1.**
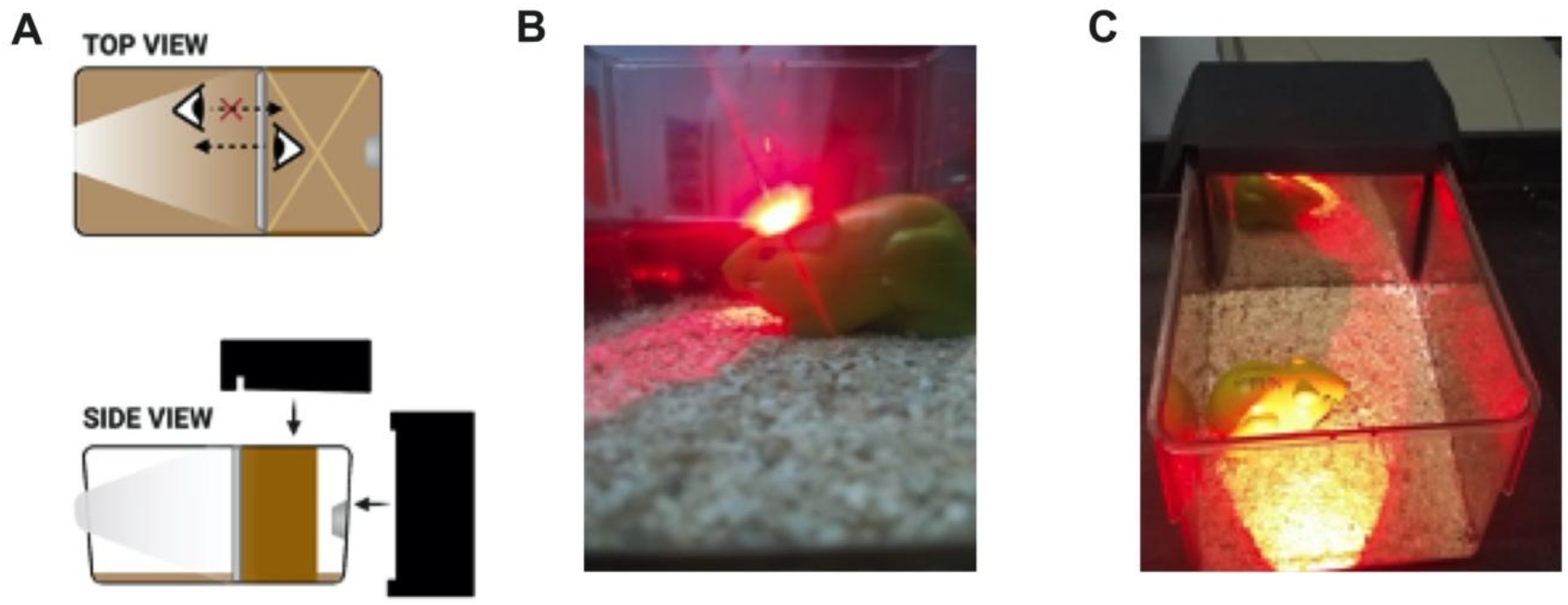
1-Way Mirror Prototype, Related to Methods. **A)** Unidirectional visual access allows the recently shocked mouse in the darkened compartment to see the cagemates on the other side of the insert, in the bright compartment. **B** and **C)** The cagemates do not see the shocked cagemate in the dark compartment but rather see their own reflections. Auditory-olfactory stimuli are exchanged over the top and sides of the 7” x 11” x 5” insert.

**Fig. S2.**
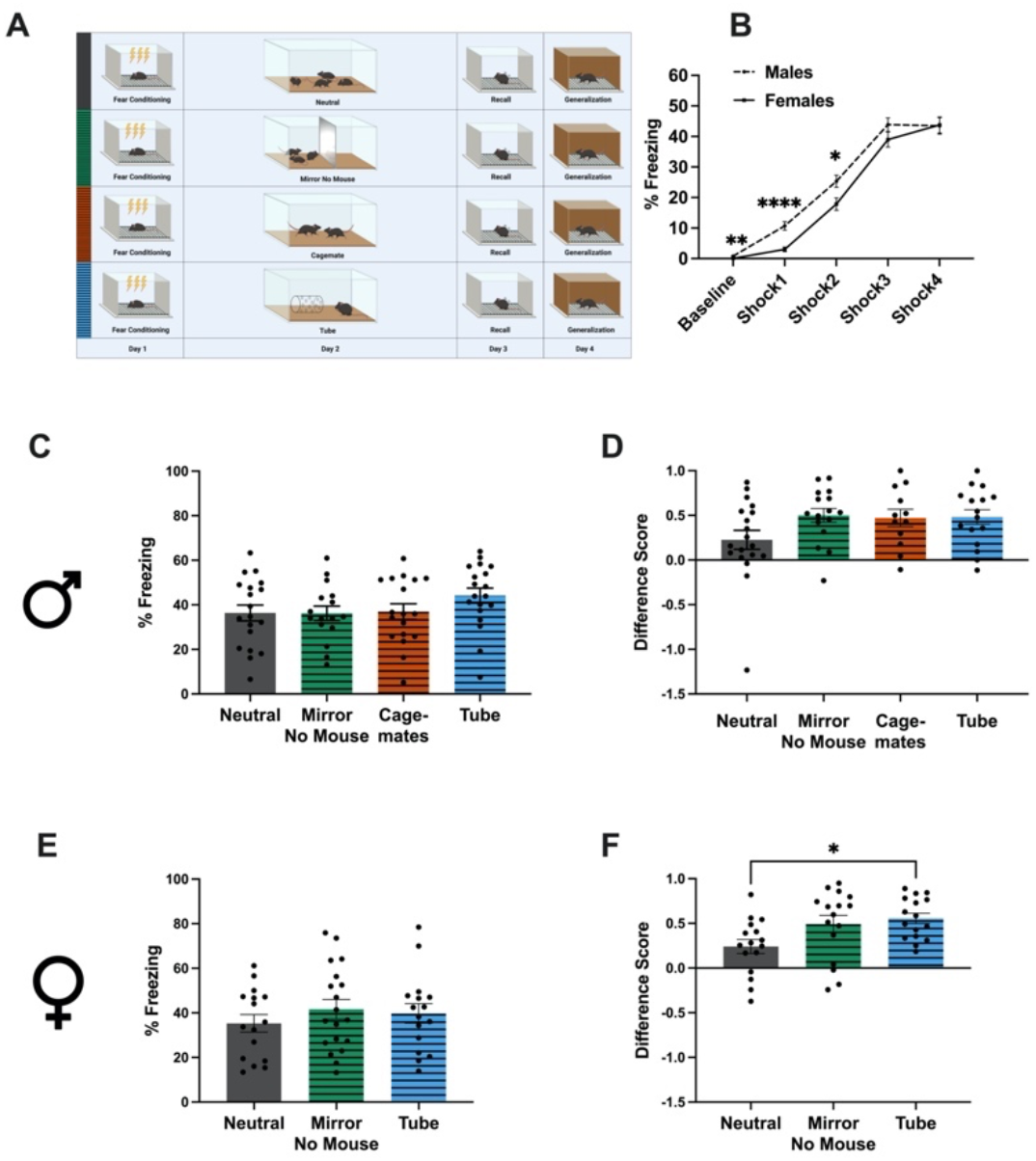
Novel objects and neutral social experiences do not impact fear recall. Related to Figure. **1.A**) Schematic representation of behavioral schedule. **B**) % freezing throughout fear acquisition in males (dashed) and females (solid). 2 Way RM ANOVA with Sidak’s multiple comparison tests of fear acquisition for males vs females, n=69 males, n=63 females; Time x Sex *F*_(4, 520)_=2.910, p=0.0212; Time *F*(2.717, 353.3)=323.8, p<0.0001; Sex *F*_(1, 130)_=4.895, p=0.0287; Subject *F*_(130, 520)_=3.594, p<0.0001; Baseline - t=3.336, p=0.0063; Shock1 - t=4.930, p<0.0001; Shock2 - t=2.626, p=0.0475; Shock3 - t=1.464, p=0.5452; Shock4 -t=0.07762, p>0.9999. **C** and **E**) Freezing levels during a 5 minute fear recall test for males (**C**) and females (**E**). Males: 1-Way ANOVA*F*_(3, 68)_=1.290, p=0.2848. Females: 1-Way ANOVA*F*_(2, 48)_=0.5864, p=05603. **D** and **F**) Differences scores, defined as [(freezing in recall test)-(freezing in generalization test)]/(freezing in recall test) for males (**D**) and females (**F**). Males: ANOVA*F*_(3, 59)_=2.172, p=0.1009. Females: Kruskal-Wallis statistic=8.115, p=0.0173, Dunn’s multiple comparisons Tube vs Neutral z=2.551, p=0.0323. Data shown as mean +/− SEM.

**Fig. S3.**
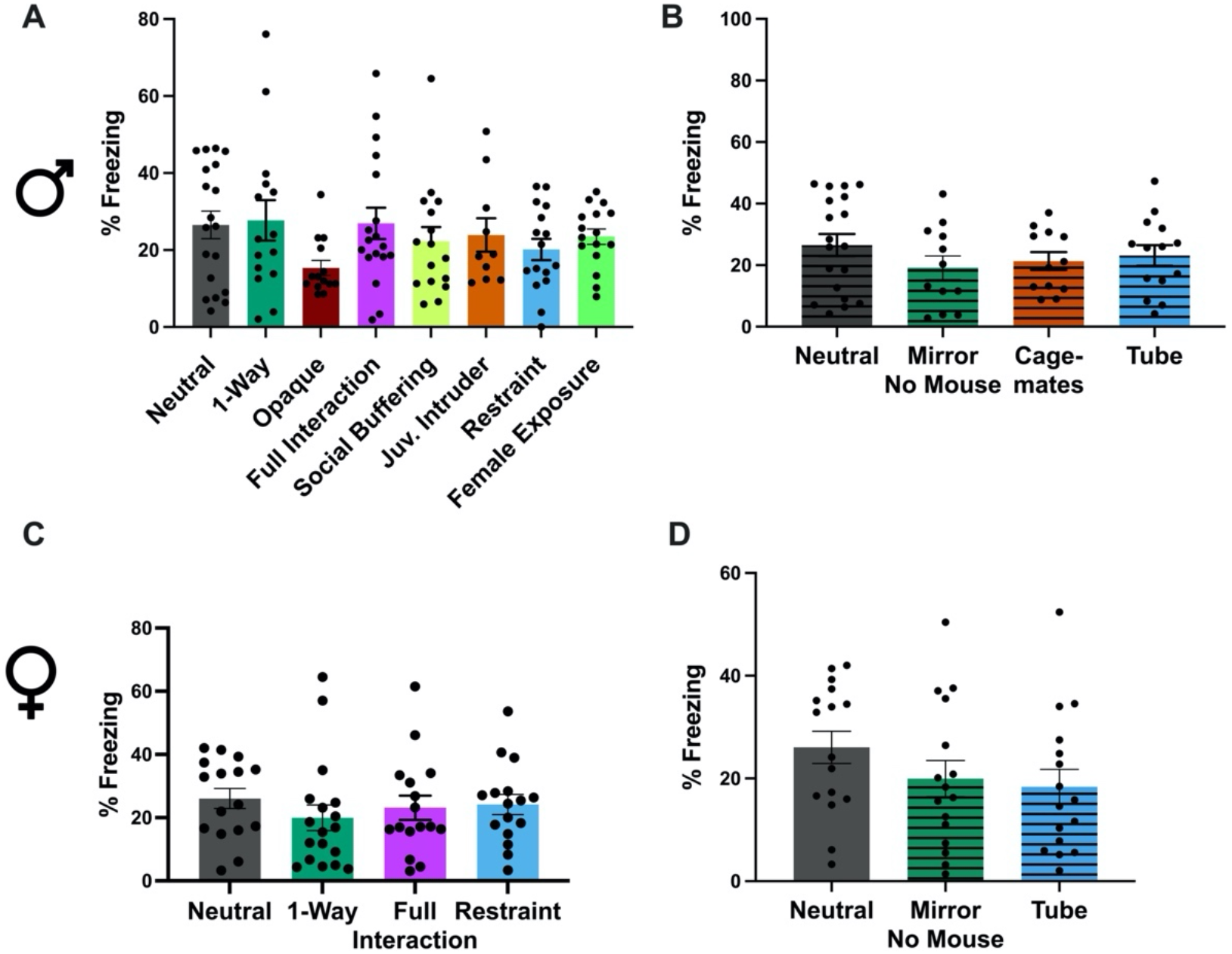
Non-normalized freezing generalization context does not differ between groups. Related to Figure 1. Each graph shows the % freezing during the generalization test, with the Neutral group included as reference. **A**) Experimental Males: Brown-Forsythe ANOVA*F*_(7, 87.07)_=1.273, p=0.2730; **B**) Control Males: 1-Way ANOVA*F*_(3, 53)_=0.8275, p=0.4846; **C**) Experimental Females: ANOVA*F*_(3, 62)_=0.5153, p=0.6733; **D**) Control Females: ANOVA*F*_(2, 45)_=1.452, p=0.2448.

**Fig. S4.**
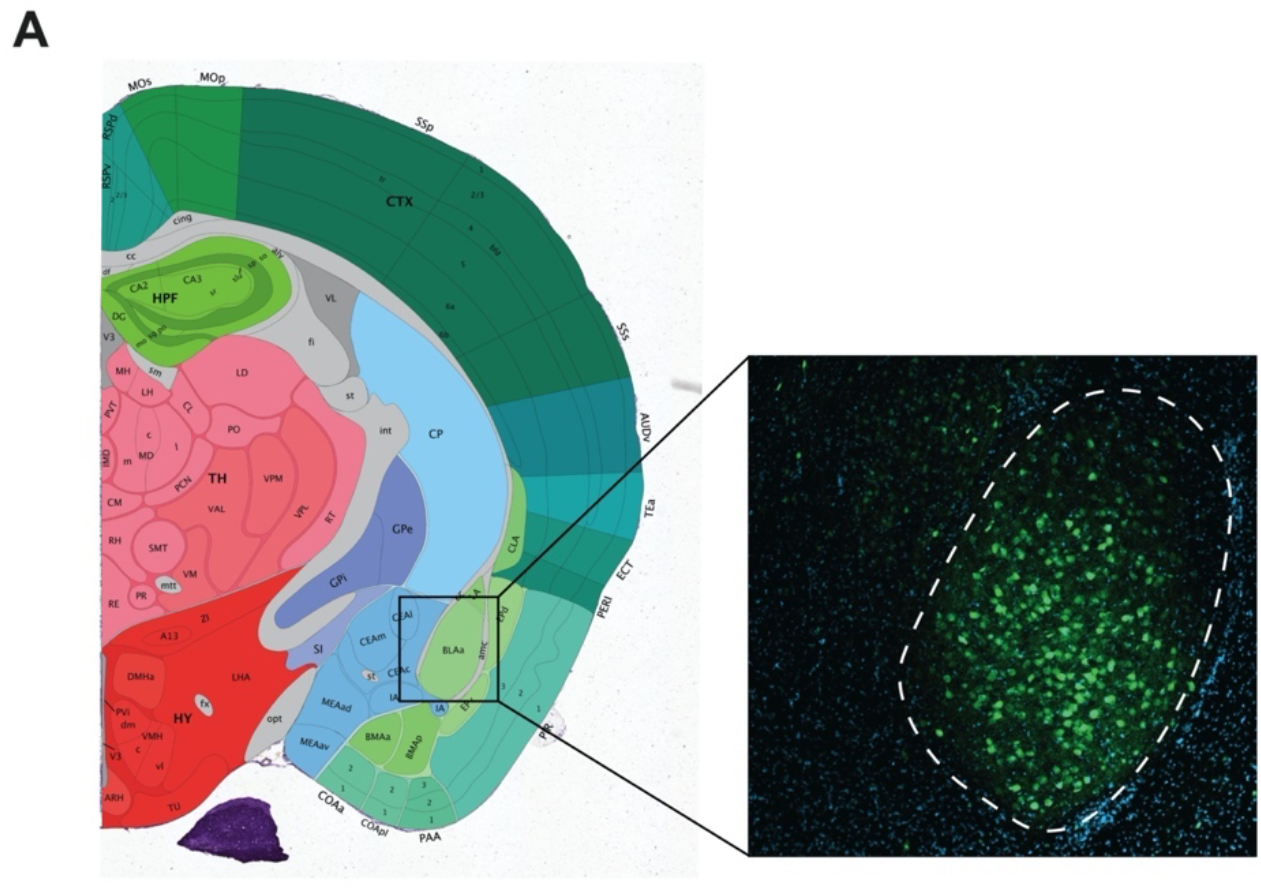
BLA histology. Related to Figure 2. **A**) Representative 10x confocal image demonstrating the bounds used to define the BLA.

**Fig. S5.**
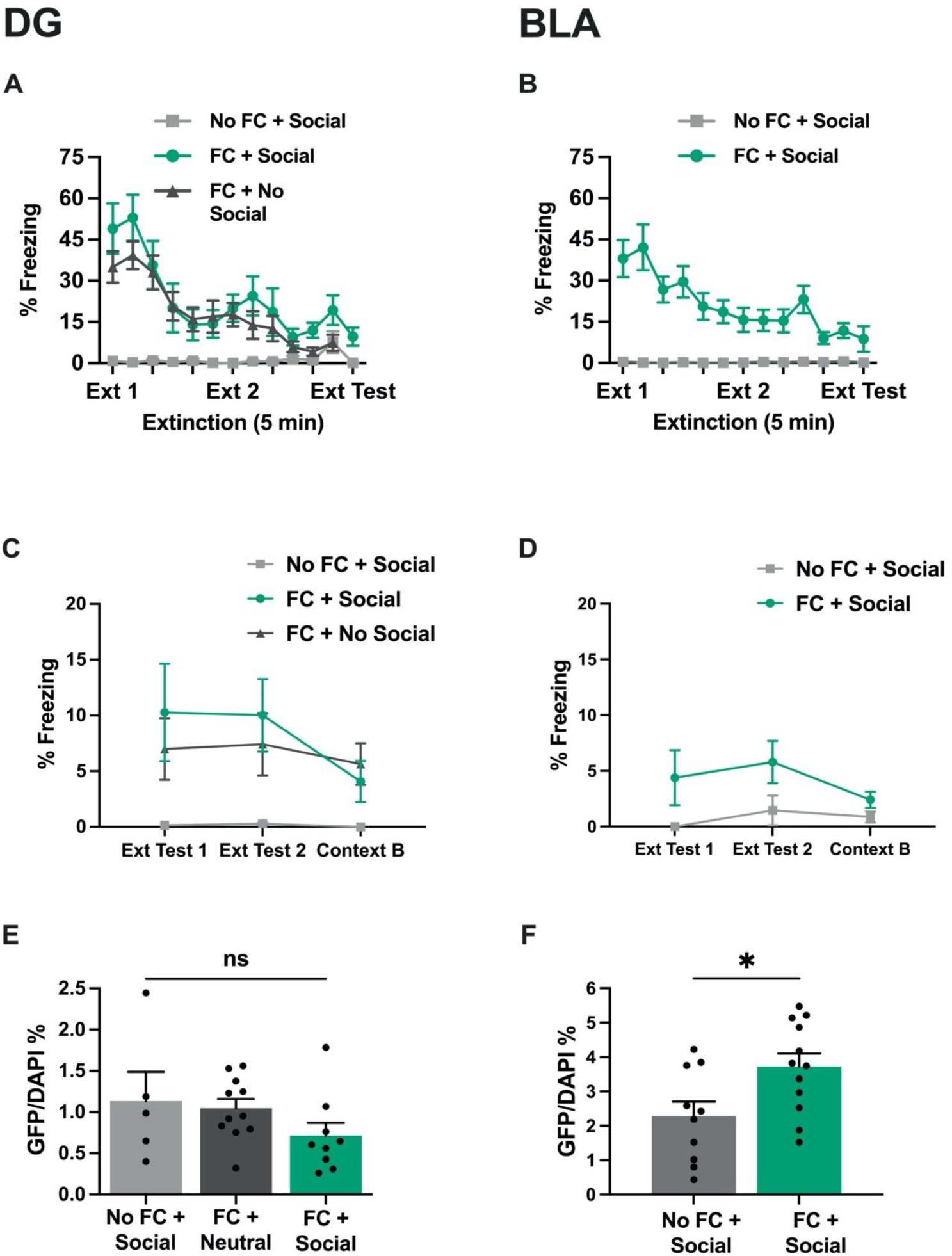
Ensemble size and extinction learning curves for optogenetically manipulated groups. Related to Figure 4. **A** and **B**) % freezing throughout extinction training across groups. Mixed-effects model (REML), **A**) time p<0.0001, group=0.0025. **B**) time p=0.0016, group <0.0001. **D** and **D**) % freezing during ext test 1, ext test 2 post stimulation, and context b. **E** and **F**) % Chr2-GFP positive cells over total DAPI positive cells. **E**) 1-Way ANOVA *F*_2, 22_=1.497, p=0.2458. **F**) unpaired 2-tailed t-test t=2.528, df=20, *p=0.02.

**Movie S1.**

Representative video of 1-way mirror paradigm for fear conditioned cagemates.

**Movie S2.**

Representative video of juvenile intruder paradigm for fear conditioned cagemates. The experimental resident mouse has two marks at the base of his tail, the intruder has a long solid line from the base to the middle of his tail.

